# Design of Ceramic Packages for Acoustically Coupled Implantable Medical Devices

**DOI:** 10.1101/836312

**Authors:** Konlin Shen, Michel M. Maharbiz

## Abstract

**Objective:** Ultrasonic acoustic power transfer is an efficient mechanism for coupling energy to millimeter and sub-millimeter implants in the body. To date, published ultrasonically powered implants have been encapsulated with thin film polymers that are susceptible to well-documented failure modes *in vivo*, including water penetration and attack by the body. As with all medical implants, packaging with ceramic or metallic materials can reduce water vapor transmission and improve biostability to provide decadal device lifetime. In this paper, we evaluate methods of coupling acoustic energy to the interior of ceramic packages.

**Methods:** The classic wave approach and modal expansion are used to obtain analytical expressions for acoustic transmission through two different package designs and these approaches are validated experimentally. A candidate package design is demonstrated using alumina packages and titanium lids, designed to be acoustically transparent.

**Results:** Bulk modes are shown to be more effective at coupling acoustic energy to a piezoelectric receiver than flexural modes. Using bulk modes, packaged motes have an overall link efficiency of roughly 10%, compared to 25% for unpackaged motes. Packaging does not have a significant effect on translational misalignment penalties, but does increase angular misalignment penalties. Passive amplitude-modulated backscatter communication is demonstrated.

**Conclusion:** Thin lids enable the use of acoustically coupled devices even with package materials of very different acoustic impedance. *Significance:* This work provides an analysis and method for designing packages that enable acoustic coupling with implantable medical devices, which could facilitate clinical translation.

## I. Introduction

Implantable medical devices (IMDs) represent a large and growing fraction of medical therapeutics and diagnostics. In the diagnostic domain, IMDs can enable not only more frequent monitoring of biometric data, but also improved spatial resolution and sensitivity compared to traditional clinical tools such as magnetic resonance imaging and computed tomography [1, 2]. Furthermore, the use of custom application specific integrated circuits (ASICs) within IMDs enables sophisticated multi-function devices for closed-loop treatment programs [3-6].

As a tradeoff, IMDs must be surgically placed in the body and thus must survive long enough *in vivo* to warrant implantation. *In vivo* lifetimes should ideally be on the scale of decades, a significant portion of a patient’s life. Premature device failure often occurs due to mechanical damage to leads and connectors, as well as corrosion and electrical shorts due to moisture penetration through packaging [7-9]. Additionally, functional devices can be rendered ineffective due to the tissue response from a foreign body, which results in device encapsulation and separation from healthy tissue, or infections due to percutaneous leads [10].

Wireless and miniaturized devices can often mitigate device failure modes arising from tissue damage caused by the implant [11-13] and micromotion effects due to stiffness mismatches [14], which may result in post-surgical hematoma, fibrosis and foreign body response. However, there is a tradeoff between energy transfer efficiency and device size. As devices get smaller, the amount of available space for energy harvesting decreases, reducing the overall amount of energy harvested by the device. With this in mind, traditional wireless power transfer modalities such as RF or inductive coupling are less effective than ultrasonic power transfer at millimeter scales, due to the improved efficiency of acoustic propagation in tissue over electromagnetic propagation [15-17]. In the past decade, acoustic power transfer has been gaining popularity as a method for powering medical devices; the number of papers discussing acoustic power transfer for IMDs has increased from roughly 6 published papers between the years 2000-2009 to almost 300 published papers between 2010-2018 (Fig. S1).

To enable further reductions in device size, passive ultrasonic backscatter communication can greatly reduce the number of components necessary on a wireless sensor [18]. In this communication scheme, implanted wireless sensors (motes) are periodically queried by a pulse of ultrasound transmitted by an external transceiver, analogous to radio frequency identification. A piezoelectric receiver converts the impinging ultrasound into electrical energy to “wake up” the mote. Information from the mote can then be encoded in the returning echo, reflected off the mote, and received by the external transceiver. This information is encoded by modulating some property of the incoming pulse, such as amplitude, frequency, or phase. In the simplest implementation of passive backscatter communication, amplitude modulation (AM) can be achieved by changing the load across the piezoelectric receiver. Ultrasonic AM-backscatter communication has previously been demonstrated as an effective method of wirelessly communicating with IMDs for electrical recordings [19, 20], stimulation [21], and temperature measurements [22].

By utilizing ultrasonic powering and backscatter communication, highly miniaturized wireless implants can be built, but these devices will require robust packaging to yield acceptable device lifetimes. Thin film polymeric encapsulants, while highly conformal, are prone to water vapor transmission, delamination, and degradation in the body environment [23]. As one example of many, Barrese et al. noted cracking in parylene-coated implants as early as 37 days *in-vivo* [7]. Bioinert ceramics and metals such as alumina, SiO2, titanium, and gold are orders of magnitude less permeable to water vapor and are also much better at withstanding aqueous saline environments [24]. To underscore this point, Fang et al. explored the moisture barrier properties of a host of materials, including polyimide, SU-8, polydimethylsiloxane (PDMS), and parylene, and found that even 100 nm of ceramic (thermally grown SiO_2_) was a superior moisture barrier to microns-thick polymer coatings [25]. Metallic and ceramic housings have been used extensively in previous work [26-29], however the combination of acoustic power transfer and ceramic or metallic packaging has not been reported on in the academic literature. Properly packaged ultrasonic backscattering implants could enable body sensor networks, in which millimeter and sub-millimeter implants report critical biometrics upon queries from external ultrasound transducer on the skin (Fig. 1).

**Fig. 1.**
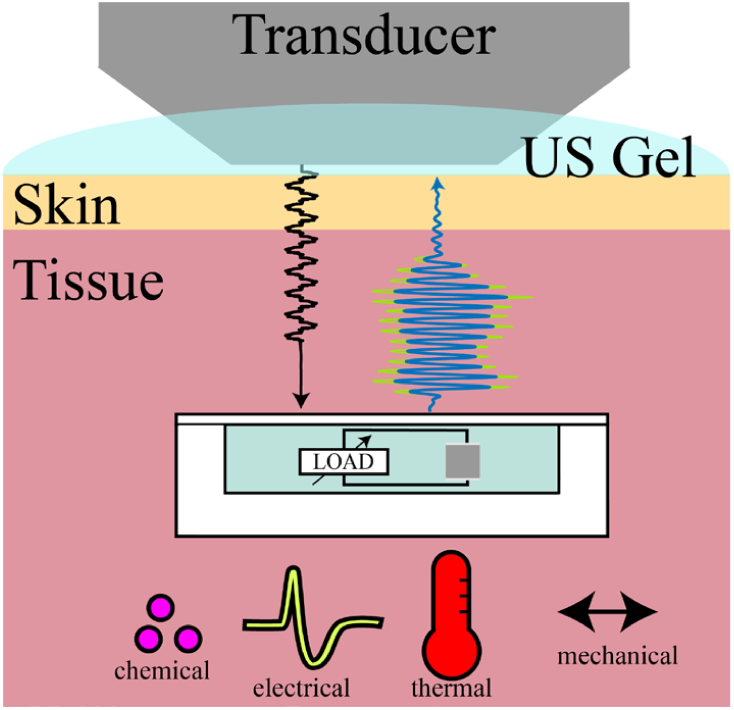
System overview for acoustically coupled implantable medical devices (IMDs) utilizing ultrasonic backscatter communication. Information from many different energy domains can be encoded in the amplitude of the backscatter (green/blue waves). For these devices to be clinically useful, packaging must be designed that both protects the devices and also allows for acoustic coupling.

In this paper, we describe the design and assembly of a ceramic package for acoustically coupled IMDs. We build and package an ultrasonically powered and backscattering sensor for electrophysiological studies and characterize packaged device performance. Finally, we expose packaged devices to a reactive accelerated aging test, characterize device lifetime, and discuss possible failure modes and solutions. A preliminary version of this work was presented in [30].

## II. Package design

As discussed above, ceramics and metals are highly desirable as package materials due to their low water vapor permeability and high biostability. However, ceramics and metals have drastically different specific acoustic impedances than that of tissue, which makes package design critical to the success of acoustically coupled IMDs. Two major strategies have emerged for coupling sound waves into packaged piezoelectric receivers. In the first strategy, the lid or a section of the lid, is thinned down to be acoustically transparent to the impinging wavelength. In the second strategy, a diaphragm is formed in the lid which has a geometry that matches the resonance frequency of the piezoelectric receiver [31] (Fig. 2).

**Fig. 2.**
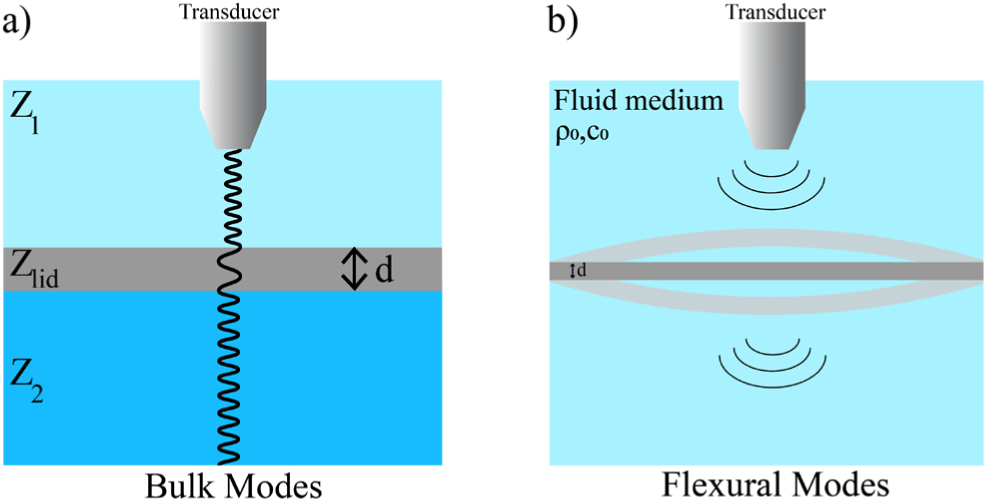
Cartoon of sound propagation through two different package lid designs. (a) Bulk mode propagation utilizes longitudinal pressure waves to transmit energy through the package material. (b) Flexural mode propagation relies on exciting resonance modes in the package, radiating acoustic energy through the medium

To determine which design is better suited for millimeter scale implants, we first consider the sound-structure interaction. When a pressure wave is incident on a solid, the resulting solid behavior is a combination of longitudinal (compression) and transverse (shear) waves propagating through the thickness of the solid. If the structure is surrounded by a fluid medium, energy can be radiated into the medium through flexural vibrations. In the thin-lid strategy, the goal is to make the solid thin enough to appear acoustically transparent. In the membrane strategy, the goal is to design the membrane in such a way to maximize radiation through the medium at the resonance frequency of the piezoelectric receiver. In the following sections, we derive and plot analytical expressions for the transmission coefficient for both coupling schemes. Model parameters can be found in Table 1.

**Table I.** Model Parameters.

| **Table I. Model Parameters** |  |
| --- | --- |
| **Longitudinal Transmission Model Parameters** |  |
| **Acoustic Impedance** |  |
| Ti ($Z_{lid}$) | 27.32 MRayl |
| Water ($Z_1$) | 1.5 MRayl |
| PDMS ($Z_2$) | 0.99 MRayl |
| Speed of Sound of Ti | 6070 m/s |
| **Flexural Transmission Model Parameters** |  |
| **Density** |  |
| Water ($\rho_0$) | 1000 kg/m<sup>3</sup> |
| Titanium | 4506 kg/m<sup>3</sup> |
| **Speed of sound** |  |
| Water ($C_0$) | 1500 m/s |
| Titanium | 6070 m/s |
| Poisson Ratio of Ti ($\nu$) | 0.31 |
| Young's Modulus of Ti | 120 GPa |
| Damping Loss Factor ($\eta$) | 0.02 |
| Plate length ($L_y$) | 2 mm |
| Plate width ($L_x$) | 1 mm |
| Plate thickness ($h$) | 0.1 mm |

### A. Bulk modes (Thin-Lid Strategy)

To model the thin-lid strategy, we assume that the primary form of energy transfer is through longitudinal waves traveling through an infinite medium with characteristic acoustic impedance *Z*_1_, going through the bulk of a finite-thickness package lid with characteristic acoustic impedance *Z*_*lid*_, and exiting into another infinite medium with characteristic acoustic impedance *Z*_2_(Fig 2a). We can calculate the transmission coefficient using the wave approach to solve for the input impedance of the combined lid and exiting medium. To do this, we will follow the derivation described by Brekhovskikh [32]. We describe a pressure wave going through the package lid at depth *z* as a combination of forward going and backwards going waves:

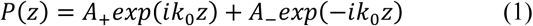

where *k*_0_ represents the wavenumber of the pressure wave in the package lid. The velocity can be calculated from 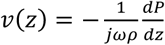 ; we obtain a velocity expression:

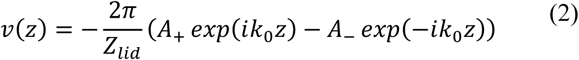

We then define the input impedance at the top interface of the package lid (including the infinite medium below it) as:

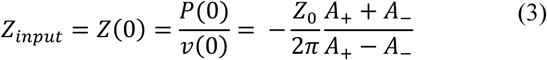

Because pressure and velocity must be continuous through the media, the interface impedance at the bottom interface of the lid, *Z*(*d*), must be identical to the characteristic impedance of the infinite medium *Z*_2_. Then:

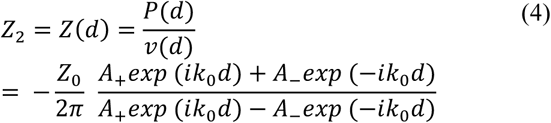

We can rearrange this equation to express *Z*_2_ as a ratio of the forward and backwards amplitudes:

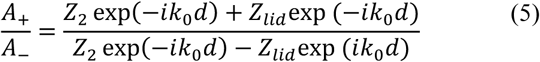

Plugging (5) into (3) we get:

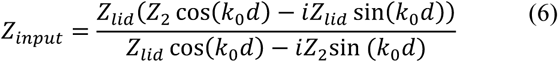

We can then determine the transmission coefficient through the whole stack as:

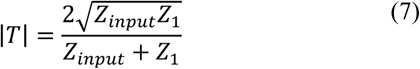

We experimentally validated this model with commercial alumina leadless chip-carrier (LCC) packages containing a piezoelectric receiver. Thin titanium sheets of varying thickness were used as package lids and the cavity was filled with PDMS to reduced flexural modes (see Methods section). The model and accompanying empirical data illustrating this phenomenon are shown in Fig. 4a and b.

**Fig. 3.**
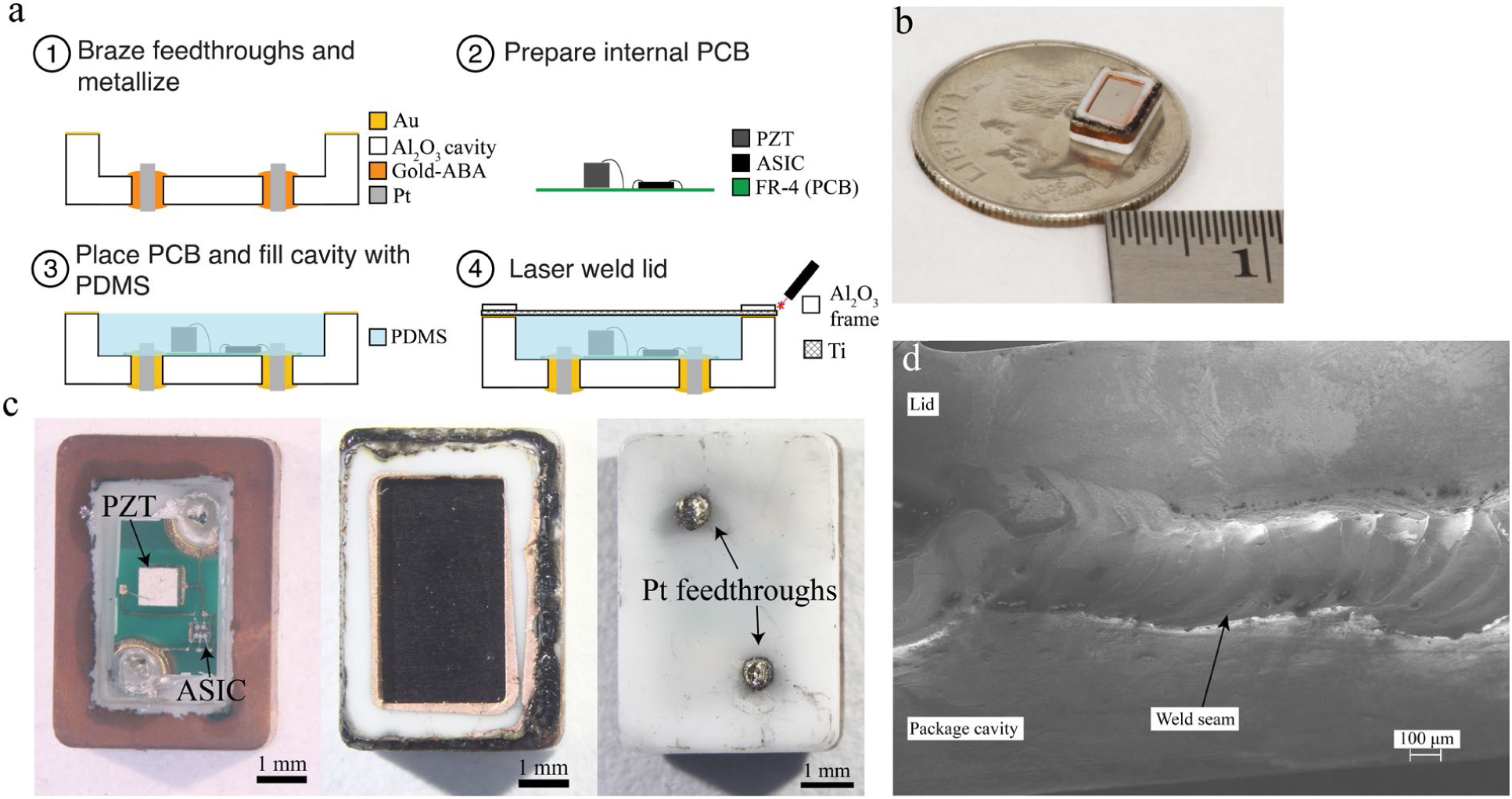
a) Packaging steps b) Picture of a sealed package on a dime. c) Micrographs of the package prior to lid sealing, post lid sealing, and from the backside. d) SEM of weld-seam where the alumina frame is welded to the package cavity. A continuous weld can be seen, joining the two pieces together.

**Fig. 4.**
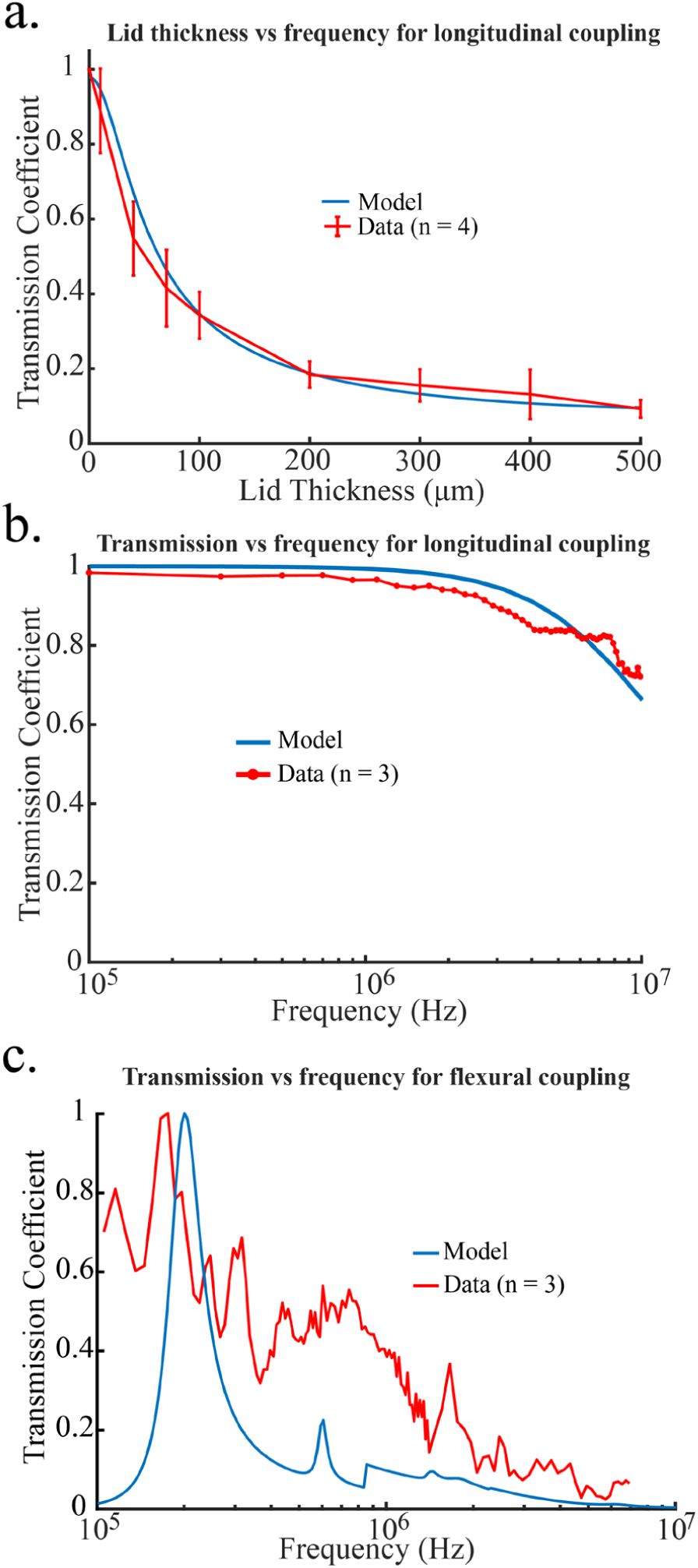
Transmission coefficients for the coupled lids. (a) transmission coefficients for longitudinal mode transmission as a function of lid thickness. PDMS is used as a backing layer. (b) Transmission coefficient for longitudinal mode transmission as a function of frequency for a 10 µm thick Ti lid. (c) Transmission coefficient for flexural mode transmission as a function of frequency for a 2 mm × 1 mm × 100 µm panel of Ti.

### B. Flexural modes (membrane strategy)

Modeling the transmission coefficient of a radiating finite plate has been studied both analytically as well as through numerical methods such as finite-element methods, boundary-element methods, and statistical energy analysis [33, 34]. While numerical methods can be more accurate than analytical methods, they are computationally expensive and time-consuming, especially at high frequencies [35]. Historically, these models were built to predict sound transmission through relatively large structural panels such as windows or building walls, in which the frequency range of interest is on the order of kHz or less. In contrast, we seek an expression for very small panels on the order of millimeters, for which resonance frequencies are on the order of 100s of kHz or higher. Thus, to reduce computational complexity, we sought an analytical expression for the transmission coefficient of a finite plate undergoing flexural vibration. Here, we loosely follow the derivation for sound transmission described by Liu et al. [36]. However, Liu’s derivation is for finite plates in air, and thus we account for fluid loading by utilizing results from Cheng et al. as well as Lomas and Hayek [37, 38]. Finally, we use the expression for radiation efficiency derived by Wallace in order to solve for sound transmission over the entire frequency spectrum [39]. Several steps of the derivation here have been omitted for brevity. A full derivation can be found in the supplement.

Let us model a rectangular titanium lid with thickness *h*, and lateral dimensions *L*_*x*_, *L*_*y*_. The incident pressure wave, *P*_*i*_, is a plane wave normal to the plate with magnitude *P*_*in*_. The equation of motion for this plate can be expressed by Kirchhoff-Love theory:

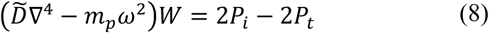

Where *W* is the normal displacement of the plate, *ω* is the angular frequency of the driving pressure wave, *P*_*t*_ is the transmitted wave, *m*_*p*_ is the area mass density of the plate given by: *m*_*p*_ = *ρ*_*p*_*h*, and 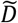 is the complex bending stiffness given by: 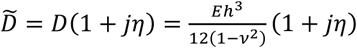, where *η* is the damping loss factor of the material. To solve for *P*_*t*_ we must know *W* which we can express as a sum of known mode shapes. For simplicity, we will assume our rectangular plate is simply supported. The mode shapes of a simply supported plate are:

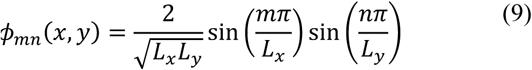

Rewriting the equation of motion with modal expansion:

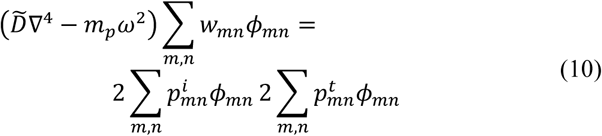

Where *w*_*mn*_, 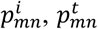 are modal coefficients. From here, we will consider each mode separately. Without loss of generality:

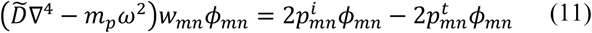

Let us define the modal wavenumber as:

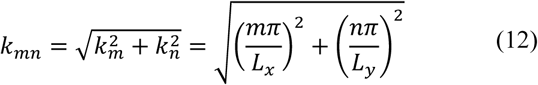

Next, let us define the modal angular frequency as:

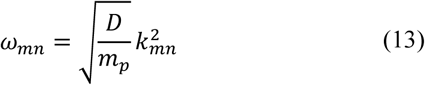

Applying the biharmonic operator to *ϕ*_*mn*_ and substituting (13) and (12) into (11) and rearranging yields:

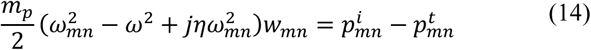

We can determine the pressure transmitted from the plate by considering the normal velocity of the plate and multiplying it by the acoustic impedance of the plate:

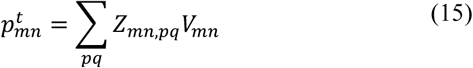

Note that here, our acoustic modal impedance is defined by two pairs of modes; this describes the effect of intermodal coupling on the pressure. The effect of mutual coupling (i.e. *mn* ≠ *pq*) has been shown to be negligible [37], so we will ignore mutual-impedance and only focus on self-impedance:

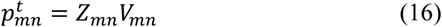

Given that *V*_*mn*_ = *jωw*_*mn*_. Plugging (16) into (14) and rearranging for *V*_*mn*_ yields:

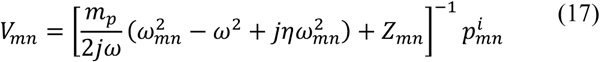

The modal impedance is a complex term:

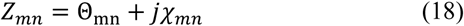

Where the real part, Θ_*mn*_, is the modal radiation efficiency and the imaginary part, *χ*_*mn*_, represents the radiation reactance, which is the result of virtual mass loading from the surrounding fluid. The modal radiation efficiency has been solved over all frequencies by Wallace and can be expressed as [39]:

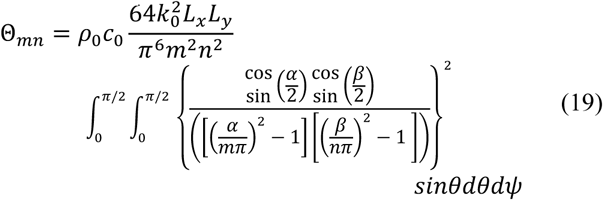

Where *α* = *k*_0_*L*_*x*_ sin(*θ*) cos (*ψ*), *β* = *k*_0_*L*_*y*_ sin(*θ*) sin (*ψ*), and the use of *cos* or *sin* in the integrand depends on whether *m* is odd (cos (*α*/2)) or even (sin (*α*/2)) and whether *n* is odd (cos (*β*/2)) or even (sin (*β*/2)).

The radiation reactance is a strong function of frequency. A complete analytical expression over all frequencies has not been determined, but an approximation used by Lomas and Hayek can be used to good accuracy [38]:

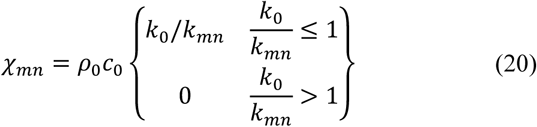

At high frequencies, the effect of water loading is negligible because at frequencies in which the phase velocity becomes supersonic (i.e. the phase velocity of the forced vibration of the plate exceeds that of the velocity of sound in water) the water is unable to vibrate in phase with the plate and thus does not add a virtual mass [33, 40].

Substituting the components of the modal impedance (18) into (17) and rearranging yields:

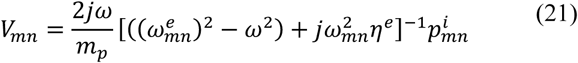

Where:

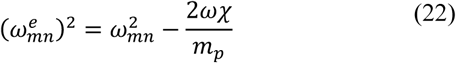

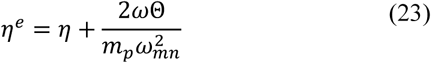

Frequently the modal amplitude of the plate velocity is expressed in terms of admittance, 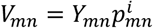, modal admittance is:

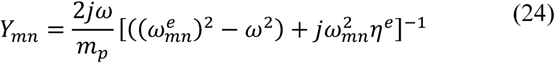

The formula for acoustic power is given by:

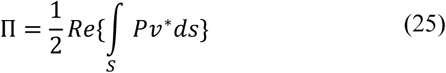

Where *S* is the area being integrated over. Then assuming a plane wave normal to the panel surface, the incident power can be given by:

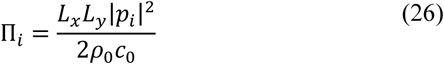

To solve for the transmitted power, we can rewrite the formula for acoustic power in terms of modes:

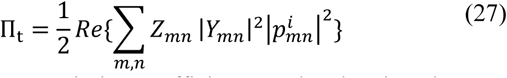

The power transmission coefficient can then be given by:

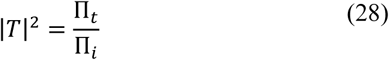

The pressure transmission coefficient can be determined by taking the square root of (28).

We validated our flexural-mode model by building titanium membranes with thin titanium sheets soldered over custom printed circuit boards (PCBs) with rectangular slots cut out to form a diaphragm. The membranes were tested in a water tank to allow for flexural vibrations (see Methods). Both the model and empirical pressure transmission coefficients are plotted as a function of frequency in Fig 4c.

## III. Methods for Model Verification, Package Assembly, and Device Testing

### A. Sound transmission model validation

Bulk-mode model validation was done by conductively epoxying a piezoelectric lead-zirconate-titanate (PZT) coupon in a leadless chip carrier (LCC, Spectrum Semiconductor, San Jose, CA) and wirebonding electrical contacts between the pads of the LCC and the piezo terminals. The LCC was then filled with PDMS to provide a backing layer for the lid to minimize flexural modes, but still allow for acoustic coupling. For lid-thickness measurements, we interrogated the piezo with continuous-wave ∼2 MHz ultrasound (corresponding to the bulk mode resonance frequency of the PZT coupon) and measured the harvested voltage while placing Ti foils (Solution Materials, Santa Clara, CA) of varying thickness over the package. For frequency measurements, we bonded a 10 µm sheet of Ti over the package with silicone and swept the frequency between 100 kHz and 10 MHz. In both cases, the transmission coefficient was determined by taking the ratio of the peak-to-peak harvested voltage at the piezo with the Ti sheet to the peak-to-peak harvested voltage at the piezo without the Ti sheet.

Flexural-mode model validation was done by building titanium membranes from 100 µm thick titanium sheet soldered to custom printed circuit boards (PCB, Bay Area Circuits, Fremont, CA). The PCBs acted as the frame of the diaphragm, with internal slots matching the diaphragm dimensions drilled through the board. A 1 cm × 1 cm pad was placed around the internal slot for the titanium to be soldered to. Due to the poor wetting of solder on the native titanium oxide, a thick copper layer (2 µm) was sputtered onto the titanium sheet. To assemble the membrane, solder paste was applied to the pad of the PCB and a flip chip bonder (finetech, Berlin, Germany) was used to align the PCB to the Ti sheet and reflow the solder.

To test the transmission through the membrane, we aligned our ultrasound transducer (Olympus, Waltham, MA) to a hydrophone (ONDA Corp, Sunnyvale, CA) and measured the received pressure of the hydrophone with and without the membrane placed in the acoustic field while sweeping the frequency, again using continuous-wave ultrasound. Transmission measurements were determined by taking the ratio of the peak-to-peak pressure at the hydrophone with the membrane to that of the peak-to-peak pressure at the hydrophone without the membrane.

### B. Package assembly

As a proof of principle, we built a ceramic package for an implantable wireless electrophysiology device using the thin-lid strategy. The size of the packaged mote is roughly 4.32 mm × 6.22 mm × 1.6 mm. Communication with the mote is achieved through passive AM-ultrasonic backscatter communication in which the recorded electrical signals modulates the load across a PZT receiver. The modulation circuit consists of a single transistor and a resistor bridge, built into a single integrated circuit (IC) [19]. The supply voltage for the transistor is provided by the PZT receiver. The IC and the piezo are mounted and electrically connected to each other on an FR-4 PCB. The PCB is packaged within an alumina housing which has a thin titanium lid to enable acoustic coupling. Platinum feedthroughs are used as electrodes to feed electrical signals recorded outside the package into the modulator for wireless reconstruction.

Material selection for these packages is important to ensure biocompatibility and prevent corrosion. Both alumina and titanium, as discussed previously, have had extensive history as biomedical implants due to their inertness and biostability. Dissimilar metals were avoided as much as possible to prevent galvanic corrosion. The metals in use (Au, Pt, Ti) were chosen due to their similar standard electrode potential, which prevents spontaneous galvanic corrosion when submerged in electrolyte [41].

The packaging of ICs and piezoelectrics imposes a relatively low temperature budget on the process. The piezoelectric properties of ceramics are endowed by asymmetry in the crystal lattice of the material. Above the Curie temperature of the material, the atomic structure experiences a ferroelectric-to-nonferroelectric transition, losing its piezoelectric properties. As a rule of thumb, processes should be held below half the Curie temperature to avoid loss of polarization [42]. For PZT, the Curie temperature is ∼350 °C, which places an upper limit on process temperatures to ∼180 °C. While other piezoelectric materials with higher Curie temperatures could be used, such as aluminum nitride (AlN) or lithium niobiate (LiNbO_3_), they have worse electromechanical coupling coefficients and there is still an upper limit on temperature set by the IC. Conventionally, ICs cannot tolerate thermal processes above 400-450 °C without permanent damage. The use of PZT rules out many conventional bonding techniques such as Au-Au thermocompression or Au-Sn eutectic bonding. In order to obtain a good seal, but not depole the piezo, we utilized laser microwelding, which delivers heat locally and thus can be used in proximity to heat-sensitive components.

The packaging process is shown schematically in Figure 3a. Custom unmetallized 99.8% purity alumina packages were 3D printed (Ceramco Inc., Center Conway, NH). First, 400 µm platinum pins were brazed into the cavity vias at 1060 °C in a vacuum environment (∼10^−5^ Torr) with a gold active-braze alloy (ABA) (Wesgo, Hayward, CA), which is an alloy composed primarily of gold (96.4%) and contains trace amounts of Ti which help the alloy wet onto the unmetallized alumina. Once the platinum pins were brazed into the cavity, the cavity was masked off with Kapton tape and a 200 nm thick gold-seal ring was evaporated onto the alumina package with a 10 nm Ti adhesion layer. This seal ring serves as a filler material during the laser welding process. Next, the internal PCB was prepared. PZT samples were cut from commercially purchased pre-metallized bulk-ceramic discs (APC 841, APC International, Mackeyville, PA) with a wafer dicing saw. The PZT and modulation IC were mounted to the PCB using silver epoxy and cured at 150 °C for 15 minutes and wirebonded to complete the circuit. The populated PCB was placed inside the package cavity and connected to the Pt-feedthroughs with silver epoxy. The package was then carefully filled with PDMS as an acoustic coupling medium. To minimize shrinkage during curing, the PDMS filled packages were cured at room temperature over 48 hours. Finally, a 10 µm sheet of titanium was bonded over the package cavity with an additional thin PDMS layer and a 3D printed alumina frame was clamped over it to provide mechanical support during laser welding. The entire ensemble was then welded together with an Nd:YAG laser (LaserStar, Riverside, RI). The device is shown in Fig. 3b,c.

### C. Package characterization

The strength of the alumina weld joint was evaluated by a tensile pull-test using a linear tensiometer (Mark-10, Copiague, NY). Two commercially available alumina LCC packages were welded together. Due to the small size of the package, handles were epoxied onto the top and bottom of the package, taking care not to get epoxy over the weld seam. The package handles were then pulled until the package broke apart while measuring the applied force. To determine the strength, the applied force was divided by area of the weld seam.

Link efficiency measurements were made by taking the ratio of the harvested power from the packaged PZT and mechanical output power of the transducer, measured by hydrophone. To calculate the harvested electrical power, we measured the harvested voltage across the piezo terminals and used:

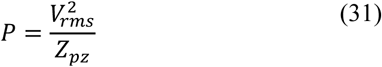

Where *Z*_*pz*_ is the electrical impedance of the piezo at resonance. Power harvesting penalty with respect to misalignment was characterized for both in-plane translation as well as out-of-plane rotation. To characterize in-plane translational misalignment, the transducer was mounted on two orthogonal manual translational stages. To characterize the rotational misalignment, the mote was mounted to a goniometer such that the axis of rotation passed through the centerline of the piezo. This was important in order to avoid possible translational offsets due to the rotation of the piezo.

To characterize the backscatter circuit, we used a sourcemeter (Keithley, Cleveland, OH) to obtain I-V curves by applying a 100 mV supply voltage while measuring the drain-source current. The gate voltage was swept from 0 to 500 mV. To determine how the amplitude of the reflected signal modulates with respect to input voltage, we swept the input voltage of the device from 0 to 500 mV in 1 mV increments while interrogating the mote with ultrasound and measuring the backscattered signal. In order to find the mote, a 300 mV, 1 Hz square wave was applied, and the transducer was steered until modulation in the backscatter signal was maximized. At each input voltage, we collected 50 backscatter pulses and averaged over them. Total modulation was determined by taking the difference between the averaged backscatter pulse at the input voltage of interest and the averaged backscatter pulse with a 0 V input. The difference signal was then rectified and a region of interest (ROI) with the largest modulation was identified. The rectified ROI in the difference signal was integrated over to provide a single value characterizing the modulation for the given input voltage. In this way, a calibration curve converting between backscatter modulation and input signal could be created. The ROI was kept constant for all input voltages during a single calibration run.

Wireless demodulation of the input signal was performed by determining the modulation value for each received waveform in the same manner as calibration and running it against the calibration curve to back out the input value.

The projected lifetime of the packaged device was determined using a reactive accelerated aging (RAA) test [43]. The devices were placed in PBS with 20 mM H_2_O_2_ to simulate the body environment after implantation. The aging set-up consisted of a jacketed flask with a flexible heater on a hot plate, a temperature controller (Omega Engineering, Norwalk, CT), and a peristaltic pump (Longer Peristaltic Pump Co., Hebei, China). The hot plate was used to provide the bulk of the heat, whereas the flexible heater was controlled by the temperature controller to provide fine control over the temperature of the flask. The pump was necessary to continuously perfuse the solution due to the short half-life of H_2_O_2_ at high temperatures. The pump speed was set to completely replace all solution in the flask in roughly 20 minutes, corresponding to the half-life of H_2_O_2_ at 90 °C (the highest temperature utilized). We monitored the performance of the device by performing device calibration daily and ended the experiment when backscatter modulation could no longer be detected. RAA was performed at both 90 °C and 80 °C. To predict the mean time to failure (MTTF) at physiological temperatures, lifetime data was fit to an Arrehnius model defined by [43]:

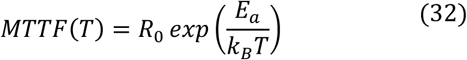

## IV. Results

### A. Bulk modes can transmit sound effectively when the lid is thin relative to the operation wavelength. Flexural modes transmit sound effectively when the resonance frequency of the mode matches the operation frequency

From the bulk-mode transmission model in (7) we see that in the limit *k*_0_*d* approaches 0, *Z*_*input*_ approaches *Z*_2_. Written another way, 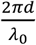, we can conclude that as the thickness of the lid becomes small relative to the interrogation wavelength, the lid becomes acoustically transparent. Note that this implies that if the medium underneath the package lid has a significantly mismatched acoustic impedance to that of the medium above the package lid, there will still be large reflections and low transmission.

Based on the empirical results in Fig. 4, we observe the following: in bulk-mode coupling, a 10 µm thick titanium lid can preserve nearly 90% of the transmitted pressure given an impinging wavelength of 2.6 mm (Fig. 3a). This corresponds to a lid thickness-to-wavelength ratio of roughly 0.003. At 1% of the wavelength, we obtain a pressure transmission coefficient of roughly 0.8. This implies that even if we were to use a 20 MHz operation frequency (approximately the bulk-mode resonance of a 100 µm PZT ceramic) a 3 µm sheet of titanium would be sufficient for 80% transmission. Furthermore, from Fig. 4b, it is apparent that this transmission coefficient is purely a function of relative thickness-to-wavelength, and that longer wavelengths preserve high transmission coefficients.

Flexural modes, on the other hand, exhibit strong transmission at a particular resonance frequency, but quickly drop off outside resonance (Fig 3c.). The construction of our membrane likely results in clamped boundary conditions rather than simply supported boundary conditions. To account for this, we implemented a frequency-dependent correction factor to predict transmission for clamped rectangular panels. For frequencies below the half the critical frequency (*f*_*c*_), which is the frequency at which the wavelength of sound in water is equivalent to the bending wavelength of the panel, the radiation power of the clamped boundary is double that of the simply-supported boundary. Between 0.5*f*_*c*_ and *f*_*c*_ the correction factor approaches unity, and for frequencies greater than *f*_*c*_ there is no difference between the clamped and simply supported panel [33, 45]. We do note that there are deviations from the modeled transmission coefficient in our measured data. The shift in the main resonance peak is likely due to sample-to-sample construction. We built a total of 4 membranes and found that while 3 of these membranes had resonance peaks slightly shifted to the left of the model, one membrane was slightly shifted to the right of the model. Furthermore, due to the size of the drill available from the commercial board house used, it was not possible to get a perfectly rectangular slot in the PCB. Finally, spurious peaks may be present in the frequency response due to the resonances in the set-up. The model assumes an infinite baffle, but the PCB was held in the set-up by an L-shaped kinematic rectangular optics mount (Thorlabs Inc, Newton, NJ), which may have contributed additional resonance peaks. Regardless, the general trend of strong transmission at a single fundamental mode and a rapid drop-off outside of resonance is still preserved.

Based on this analysis, we can make the conclusion that while relying on flexural modes will optimize transmission, we can still obtain high (< 0.8) transmission coefficients with bulk-modes. Since the bulk-mode response is more lenient with respect to frequency than flexural-modes, building packages based on bulk-modes may be more effective than packages based on flexural-modes. It is worth pointing out that the piezoelectric efficiency is higher when the medium surrounding it is fluid rather than solid due to damping of the piezo. The effect of PDMS encapsulation of the piezo results in roughly a 50% decrease in voltage harvest compared to a bare piezo (Fig. S2). Thus, while bulk-modes enable a wider bandwidth, flexural-modes may be more appropriate for applications requiring higher harvested powers.

### B. Alumina lids and package cavities can be joined with laser welding

Fig. 3d shows an SEM image of the weld seam. The scalloped features show the overlap of the pulses and the seam shows that the two pieces of ceramic have fused into a single piece rather than melting the gold-seal ring as a filler. During tensile pull tests, out of the three tested weld seams, we were only able to separate one package. The measured bond strength was roughly 26 MPa. In the other two packages, the handles broke off prior to the weld breaking, implying that the weld strength is at least 8 MPa. These values are similar to those of Au-Si eutectic bonding [46], indicating that the laser welded lids are robust.

### C. Packaging reduces power harvesting efficiency and penalizes rotational misalignment more heavily

Figure 5 shows the effects of the packaging on power harvesting and misalignment penalty compared to an unpackaged device. The unpackaged mote has a link efficiency of roughly 25% when perfectly aligned; the packaged mote efficiency is roughly 10% with perfect alignment. Thus, the mean penalty in power harvest, as shown in Fig. 5a., due to packaging is roughly 40%. The effect of packaging on translational misalignment is minimal (Fig. 5c), but due to the walls of the package, there is a substantial penalty due to angular misalignment for the packaged mote as opposed to the unpackaged mote (Fig. 5b). We note that the 3 dB point for the unpackaged mote occurs when roughly 20 degrees off axis, while the 3 dB point for the packaged mote occurs roughly 10 degrees off-axis.

**Fig. 5.**
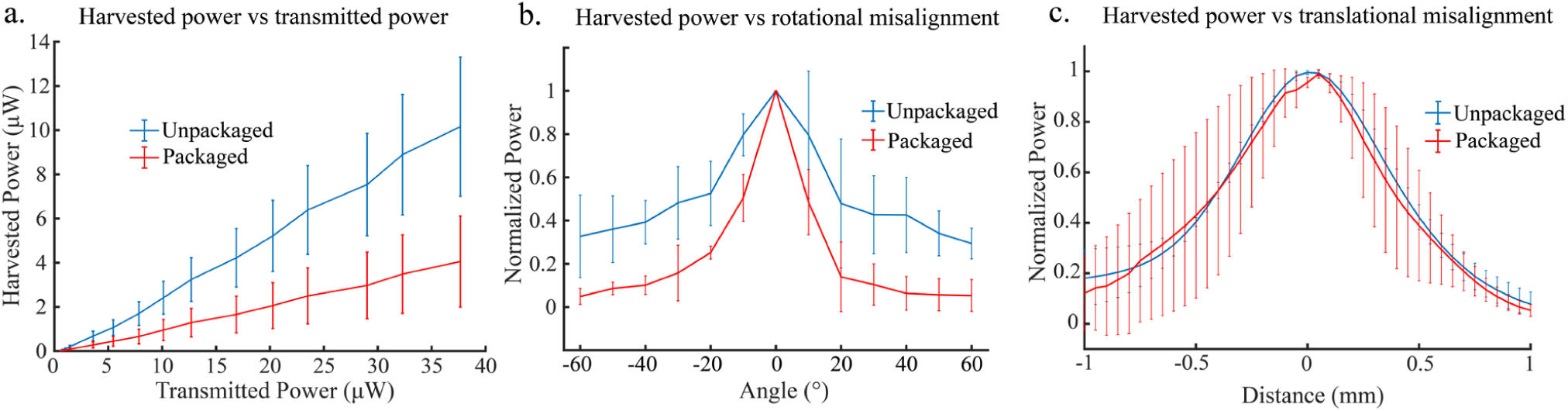
Power harvesting performance of packaged mote with unpackaged mote as a control. a) Received power as a function of transducer amplitude. b) Angular misalignment effect on power harvest. c) Effect of translational misalignment on power harvest

### D. Wireless ultrasonic backscatter communication can be performed through ceramic packages

Figure 6a shows the calibration curves used to demodulate input signals from ultrasonic backscatter plotted against the IV characteristics of the modulation circuit. There is good agreement between the IV behavior of the modulation circuit and the backscatter modulation, demonstrating that amplitude-modulated backscatter communication can be used with the packaged devices. Figure 6b shows that we are capable of wirelessly reconstructing input signals of varying amplitude and frequency with good fidelity (correlation coefficient R = 0.88). Here, we input a varying square wave pattern consisting of 1 Hz 500 mV square waves, 2 Hz 500 mV square waves, 1 Hz 450 mV square waves, and 2 Hz 400 mV square waves.

**Fig. 6.**
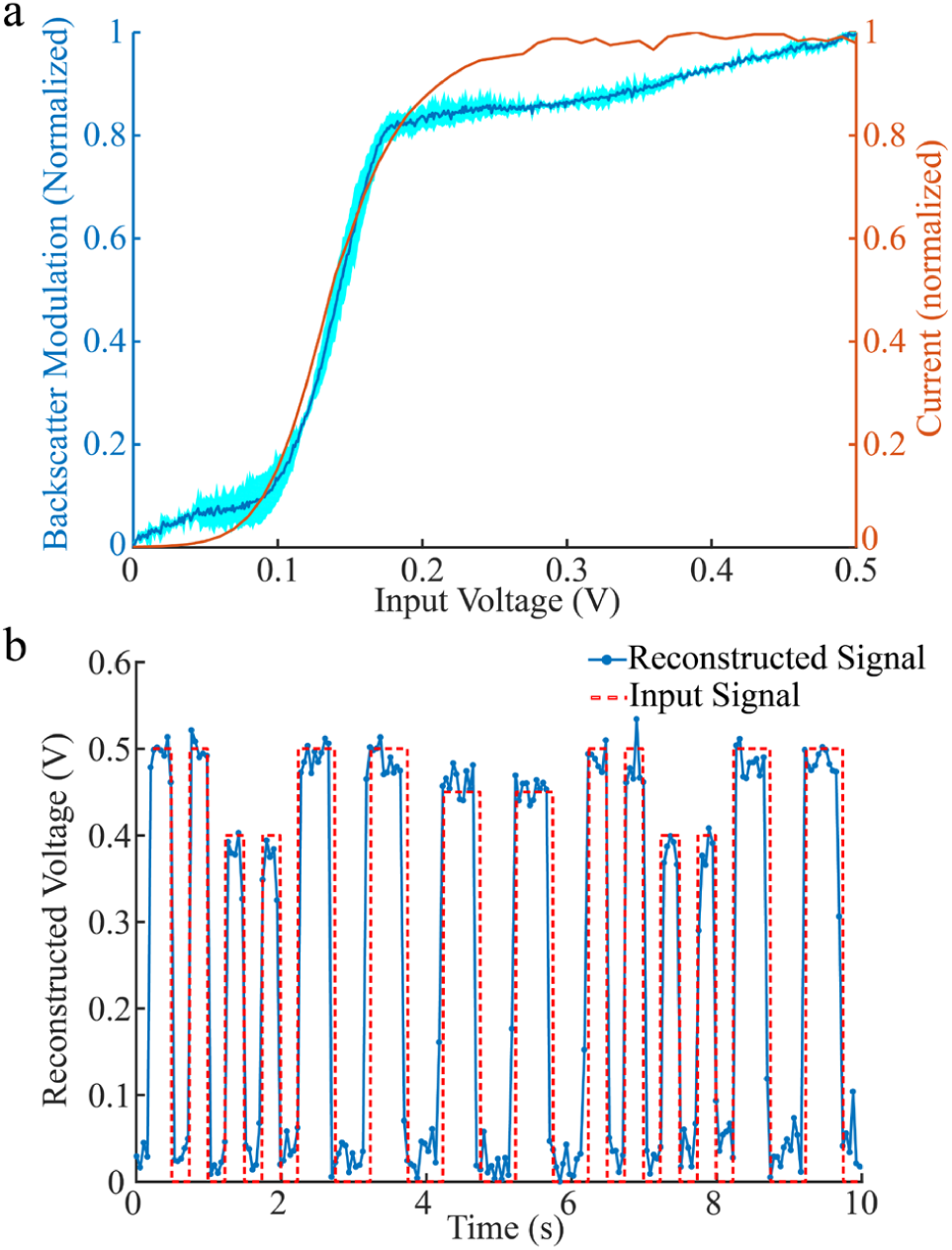
Demonstration of wireless backscatter communication in packaged motes. a) Backscatter modulation shows a good agreement with the IV characteristics of the modulation circuit (dark blue indicates the mean, light blue indicates the standard deviation, n = 3). b) The blue traces show the signal demodulated from the backscatter, which is in agreement with the input signal (red). As shown, signals of various frequency and amplitude can be reconstructed.

### E. Packaged motes have a predicted lifetime around 5 months

Figure 7 shows an example reconstruction from an aged device at 90 °C for 3 days. The reconstruction is still faithful to the input signal, with a correlation coefficient of 0.95. We found that devices failed after aging for 4 days at 90 °C and 8 days at 80 °C. Fitting this to (32), we predict a lifetime at 37.5 °C to be roughly 158 days.

**Fig. 7.**
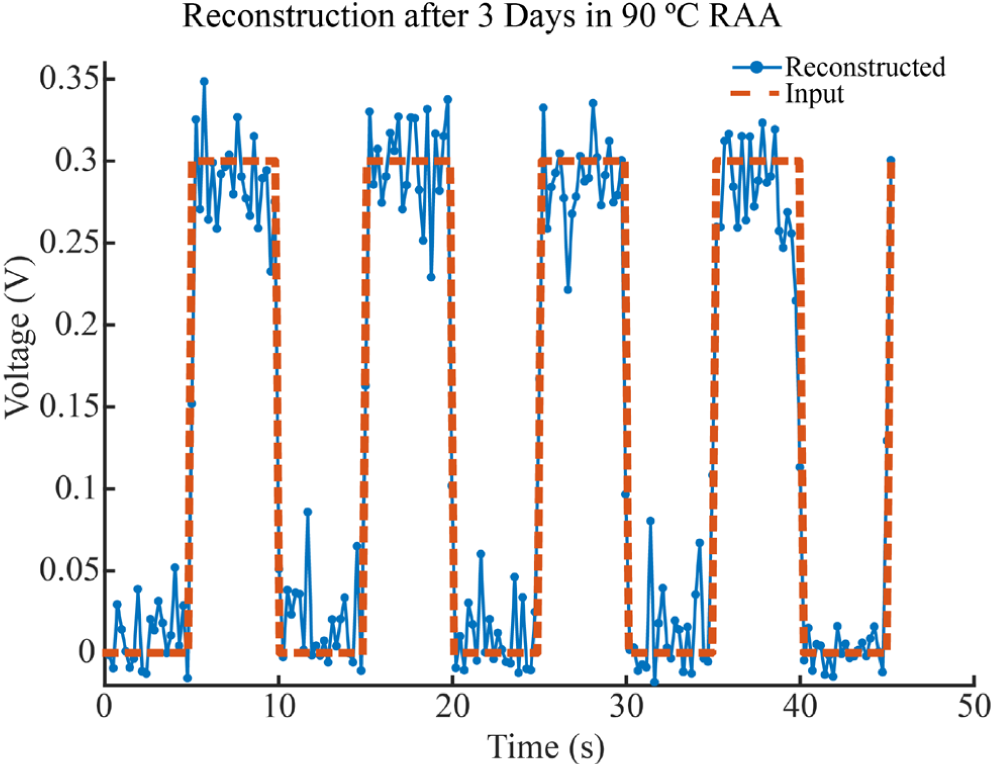
Wireless reconstruction of 100 mHz, 300 mV square waves after 3 days of RAA at 90 C and 20 mM H_2_O_2_ concentration demonstrating that the device can continue to operate and communicate after aging.

## V. Discussion

### A. Ceramic packages with thin titanium lids can be used for packaging ultrasonically coupled devices

Our results show that thin Ti lids and laser welding can enable ceramic and metal packaging of ultrasonically coupled implantable devices. By utilizing bulk modes in the 10 µm Ti sheet induced by 2 MHz ultrasound, we were able to attain 10% link efficiency as well as effective backscatter communication. While our ultrasound transmission model and experimental data suggest higher efficiency transmission with flexural modes, we chose to utilize bulk modes to simplify lid design since the resonant frequency of the lid needs to match the resonant frequency of the piezo in order to allow efficient transmission using flexural-mode coupling. Standard microfabrication methods can be used to build lids with acoustic windows that resonate at the frequency of the packaged piezoelectric receivers, and that would also allow for higher efficiency voltage harvesting since the piezo could be free to vibrate in fluid. Regardless, we find that using bulk-mode transmission is sufficient for amplitude-modulated ultrasonic backscatter communication.

### B. Microcracks in the package weld seam compromise the hermeticity of the package

We project our packaged devices to be capable of lasting within the body for several months; this lifetime is similar to that of conventional polymer-based encapsulation techniques [25, 47-49]. Although we observed complete joining of the alumina frame to its package, in some places of the weld seam, there were micro-cracks with widths up to 10 µm (Fig. S3), which likely served as moisture ingress points. Thus, we believe that the main moisture barrier here is not the housing but rather the PDMS and improving the seam quality would greatly increase the hermiticity and thus lifetime of the package.

To improve the hermiticity of the weld seam, a thicker filler material could be used to join the pieces so that the laser welding process is more akin to a laser brazing process. The filler could be anything from glass frit to an active-braze alloy or a eutectic paste [50]. Previous work by Lichter et al. showed that diamond capsules could be hermetically laser-brazed together using gold-ABA [51]. For pure ceramic-to-ceramic bonding, custom laser welding systems utilizing non-linear optics could also be employed. Watanabe et al. demonstrated joining of glasses with drastically dissimilar thermal expansion coefficients using ultrashort (femtosecond) laser pulses without the need of a filler material [52]. Itoh and Ozeki demonstrated hermetic sealing of alumina to borosilicate glass using a similar system [53].

Finally, ceramic-to-metal welds could be utilized instead of ceramic-to-ceramic welding. This method of welding was utilized in the clinically tested Bion microstimulator [28], which sought to build a wireless RF based implantable medical device with lifetimes greater than 10 years. In this device, a tantalum electrode was laser welded to a borosilicate glass capsule using a dual CO_2_ and Nd:YAG laser, resulting in a hermetic seal [54]. Thus, while in our work, we were not able to obtain a laser welded hermetic seal, we believe that weld optimization is a straightforward task.

### C. Alternative lead-free piezoelectric materials are compatible with our packages

In this work, we used PZT as our piezoelectric material due to its high elecromechanical coupling coefficient. However, the use of lead-based piezoelectrics in implantables is undesirable due to the cytotoxicity of lead. Barium titanate (BaTiO_3_) is a promising candidate to replace lead-based piezoceramics as it has a higher electromechanical coupling coefficient than other lead-free piezoceramics such as LiNbO_3_ and AlN [55]. Switching to BaTiO_3_ would be 70% as efficient as PZT (Fig. S4), which is acceptable for our use-cases. The drawback to using BaTiO_3_ is its low curie temperature (120 °C), which sets an even more stringent temperature budget. However, room temperature curing conductive epoxies could be used, and we do not believe that the laser welding process currently employed would depole the piezo.

### D. 3D printed packages and wedge wirebonding limits the extent of miniaturization

Using the described packaging scheme, we were able to build slightly smaller packages with dimensions 5 mm × 4 mm × 1.6 mm. The major constraint on miniaturization is the size of the internal PCB. The IC and PZT coupon used in this work together occupy about 1 mm^2^. Additional space is required for wirebonding target pads, interconnects, and empty space to accommodate the wirebonder head. As a result, internal PCBs must have dimensions on the order of millimeters. The 3D printing process has a minimum wall thickness of 120 µm, so the overall dimensions of the package would be ∼0.25 mm larger in each dimension than the PCB. Sub-millimeter packages are not easy to build using this packaging method but could be realized utilizing flip-chip processes and standard microfabrication techniques.

## VI. Conclusion

In this paper, we evaluated two different strategies for coupling ultrasound into ceramic/metallic packages. We determined analytically that bulk modes are more effective at transmitting acoustic energy into packages than relying on radiated sound energy and verified that claim experimentally. We then built proof-of-concept alumina packages with thin titanium lids utilizing a laser welding process to keep the heat transferred to the piezo at a minimum. We demonstrated that our packaged devices can both harvest ultrasonic energy as well as communicate information via amplitude-modulated ultrasonic backscattering. While the package is not significantly more effective in protecting the electronics from a corrosive environment than traditional polymer encapsulations, this analysis and fabrication method provide guidelines for developing packaging for acoustically coupled implants and similar packaging strategies have been used to build hermetic capsules for IMDs [51, 56].

Acoustics have been steadily gaining momentum as an efficient way to couple energy into deeply implanted or mm to sub-mm scale implantable medical devices. While the demonstrated device records electrophysiological potentials, other ultrasonically coupled devices for different applications have been shown such as stimulation, temperature sensing, and oxygen sensing. Thus, this analysis of acoustic coupling packages and assembly method demonstrates a path towards long term viability and clinical translation of acoustically coupled IMDs.

## Acknowledgment

The authors would like to thank David K. Piech for contributions to analysis software and discussion, Jerry Walias of Inta Technology for brazing and laser welding, and Soner Sonmezgolu, Kyoungtae Lee, and Oliver Chen for valuable discussion. Metal deposition and dicing steps were performed in the Marvell Nanofabrication Laboratory at UC Berkeley. M.M.M. is a member of iota Biosciences, inc.

