## Supplemental Materials for "Design of Ceramic Packages for Acoustically Coupled Implantable Medical Devices"

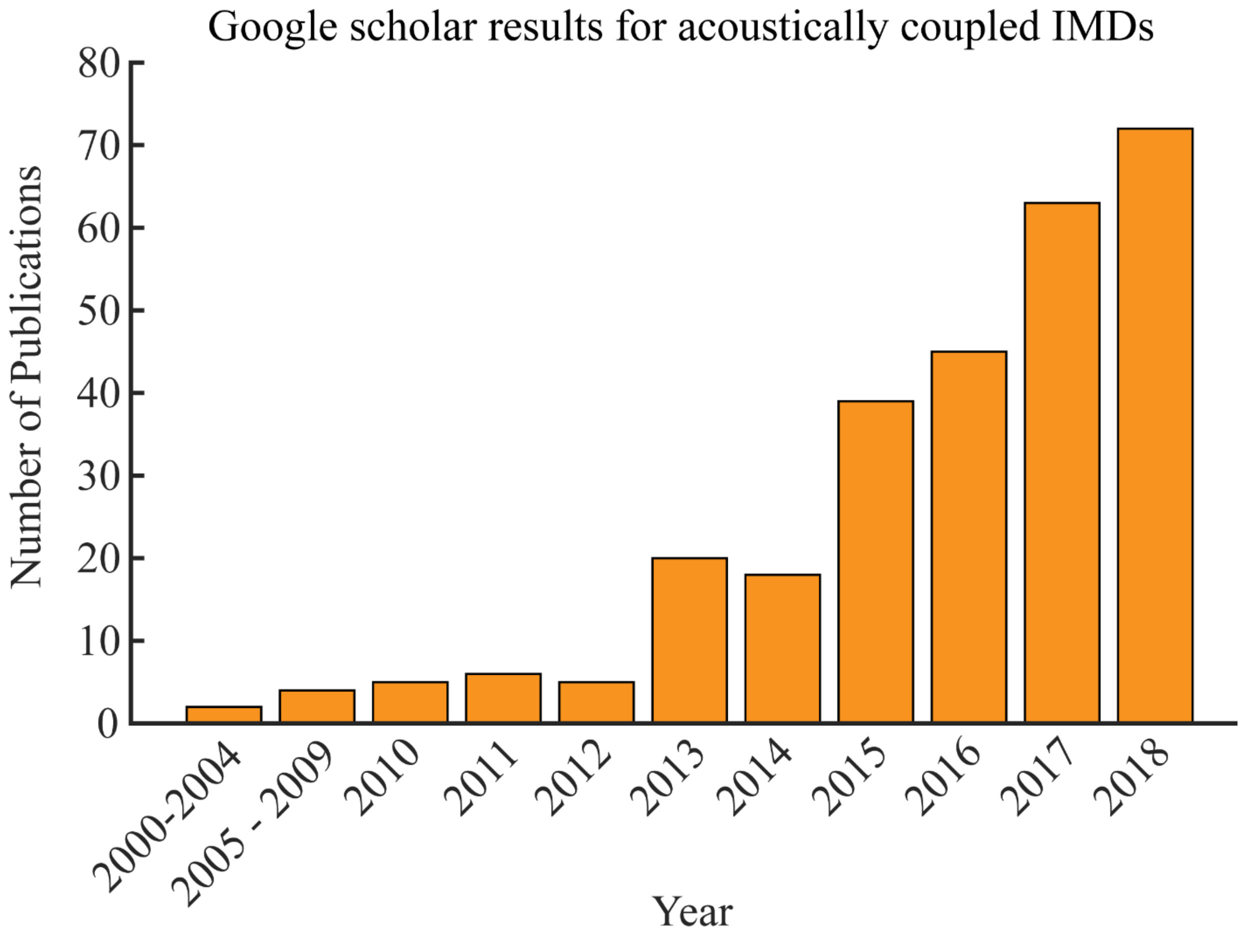


**Supplemental Figure 1**. Trend in the number of published academic papers on acoustically coupled implantable medical devices (IMDs) over the past two decades. Interest in the use of acoustics for wireless energy transfer has increased significantly since the early 2000s. Results were pulled from google scholar results using the keywords: wireless implantable medical device "acoustic power transfer" OR "ultrasonic power transfer" OR "ultrasonic energy harvesting" OR "acoustic energy harvesting"

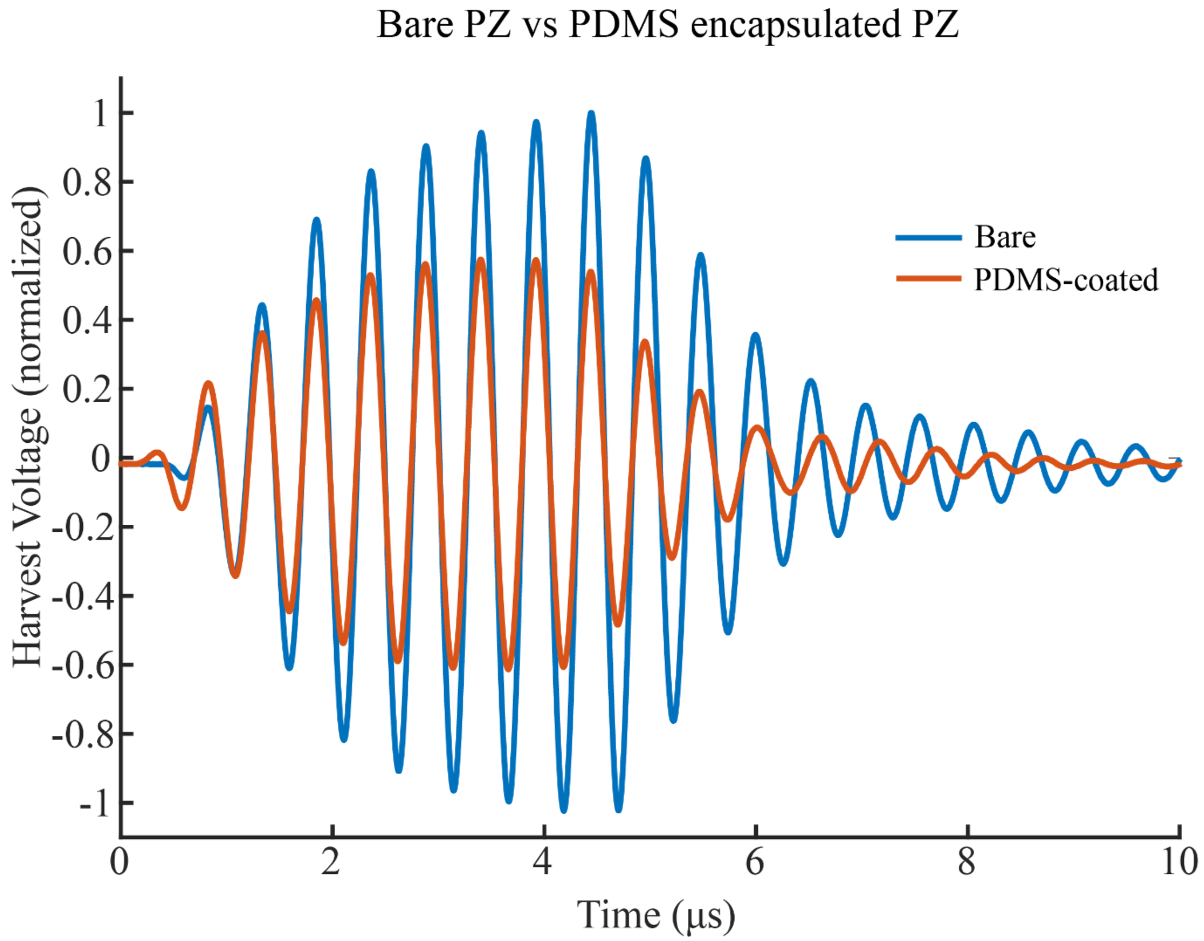


**Supplemental Figure 2.** PZT coupons potted in PDMS are significantly more damped than bare piezos. The reduction in harvested voltage due to PDMS encapsulation is roughly 50%. Both bare and PDMS-coated piezos were mounted on FR-4 PCBs and tested with an 8-cycle pulse-train at roughly 2 MHz in castor oil.

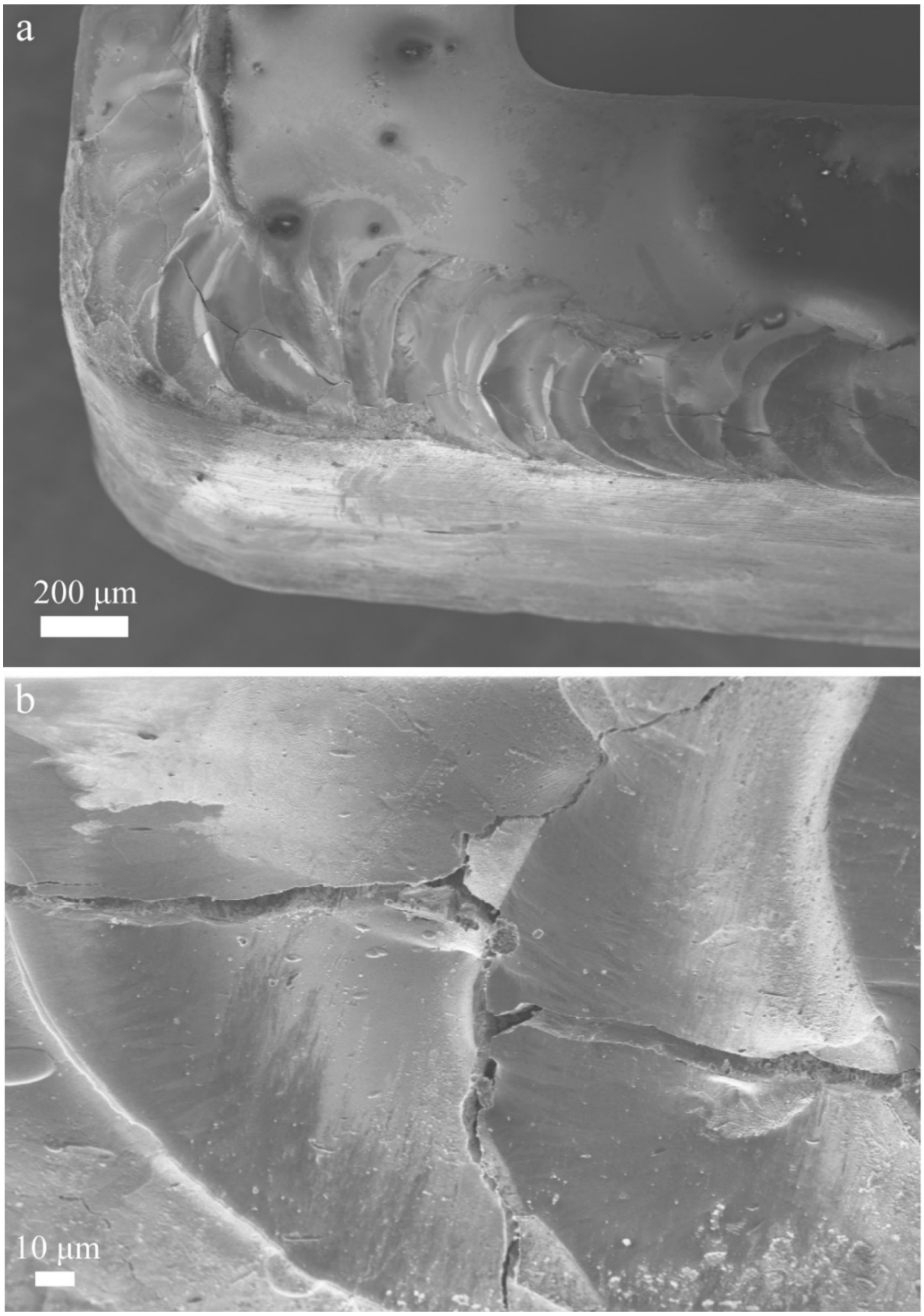


**Supplemental Figure 3.** SEM images of micro-cracks in the weld seam due to the thermal stress generated by temperature gradients near the weld seam. These cracks can be as wide as 10 µm and compromise hermiticity. Various methods can be used to counter crack formation such as using a filler material to perform laser brazing [1], local preheating of the weld seam using a second laser to minimize thermal gradients [2], or using ultrashort laser pulses [3].

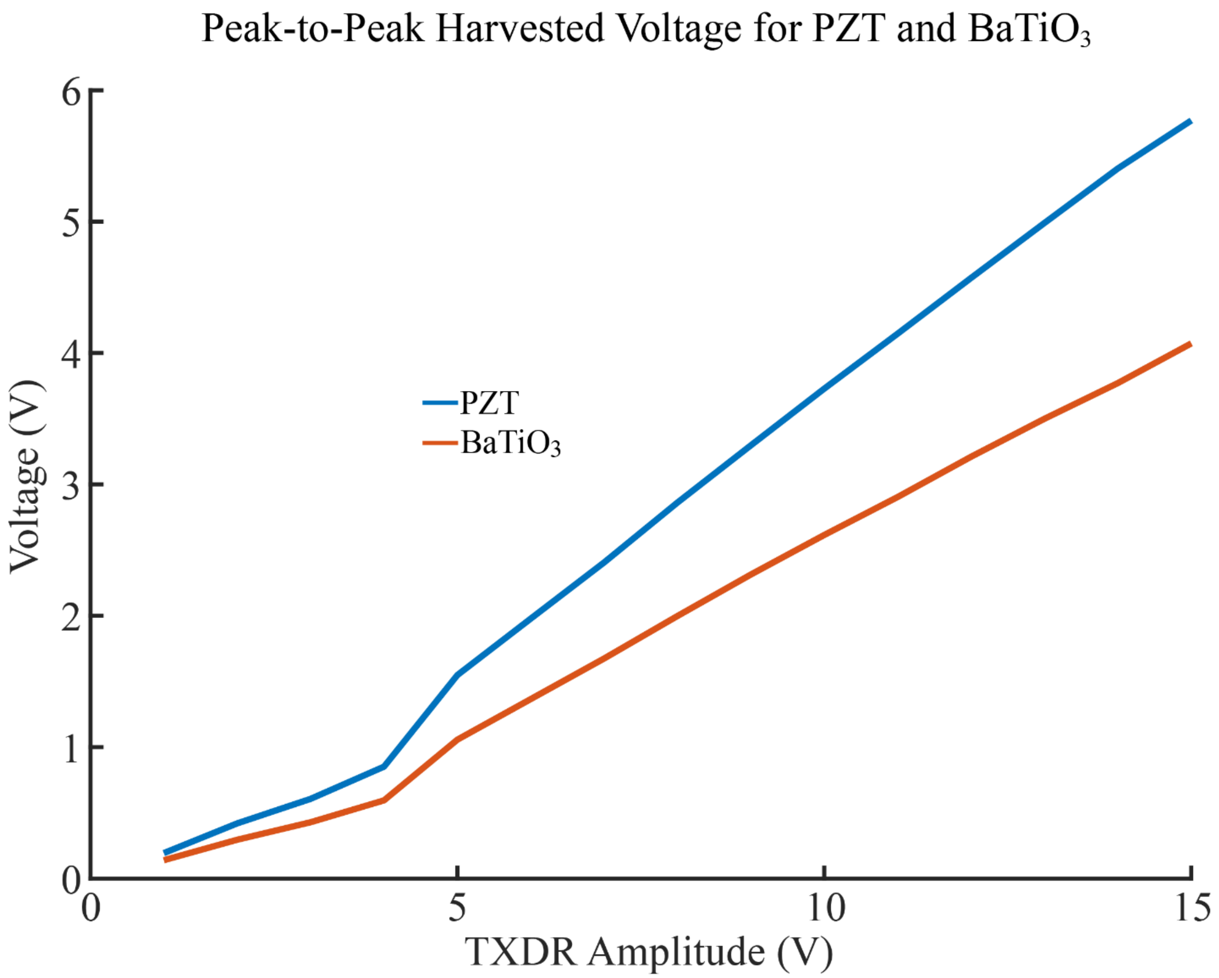


**Supplemental Figure 4**. Comparison of voltage harvesting between $BaTiO_3$ and PZT. $BaTiO_3$ is a lead-free and biocompatible piezoelectric material [4]. Peak voltage harvest for $BaTiO_3$ is roughly 70% that of PZT which is sufficient for powering our passive amplitude-modulated backscatter circuit. PZT and $BaTiO_3$ samples were mounted on polyimide PCBs and tested with an 8-cycle pulse train at the resonance frequencies of the samples (~2 MHz and ~2.5 MHz for PZT and $BaTiO_3$ respectively) in castor oil.

**Supplementary Section 1: Full derivation of the transmission coefficient for a finite titanium plate in water**

In this derivation, we utilize the modal approach, following the derivation done by Liu et al. [5]. However, their derivation examines a spherical wave propagating through air; in our case, we instead consider a plane wave propagating through a liquid (water) medium. Because the liquid medium is of similar density to the plate ($\rho_{water} = 1000\ kg/m^3,\ \ \rho_{Ti} = 4506\ kg/m^3$), we cannot ignore the effect of fluid loading on the plate. To take this into account, we utilize results from Cheng et al. [6] and Lomas and Hayek [7].

Assume a simply supported plate with thickness $h$, and lateral dimensions $L_x, L_y$. The incident pressure wave, $P_i$, is a plane wave normal to the plate with magnitude $P_{in}$. The equation of motion for this plate can be expressed by Kirchhoff-Love theory:

$$\left(\widetilde{D}\nabla^4 - m_p\omega^2\right)W = 2P_i - 2P_t$$

Where:
$W$ is the normal displacement of the plate
$\omega$ is the angular frequency of the driving pressure wave
$P_t$ is the transmitted wave
$m_p$ is the area mass density of the plate given by: $m_p = \rho_p h$
$\widetilde{D}$ is the complex bending stiffness given by: $\widetilde{D} = D(1 + j\eta) = \frac{Eh^3}{12(1-\nu^2)}(1 + j\eta)$, where $\eta$ is the damping loss factor (intrinsic to material). In this derivation, we assume the damping loss factor for Ti is similar to that of Al ($\eta = 0.02$).

We can use modal expansion to express the displacement, incident pressure, and transmitted pressure as a sum of mode shapes. The mode shapes of a simply supported plate are given by:

$$\phi_{mn}(x, y) = \frac{2}{\sqrt{L_x L_y}} \sin\left(\frac{m\pi}{L_x}\right) \sin\left(\frac{n\pi}{L_y}\right)$$

Then expanding the displacement, incident pressure, and transmitted pressure:

$$W(x, y) = \sum_{m,n} w_{mn}\phi_{mn}(x, y)$$

$$P_i(x, y) = \sum_{m,n} p_{mn}^i \phi_{mn}(x, y)$$

$$P_t(x, y) = \sum_{m,n} p_{mn}^t \phi_{mn}(x, y)$$

Where $w_{mn}$, $p_{mn}^i$, $p_{mn}^t$ are modal coefficients.

Rewriting the equation of motion with modal expansion:

$$\left(\widetilde{D}\nabla^4 - m_p\omega^2\right)\sum_{m,n} w_{mn}\phi_{mn} = 2\sum_{m,n} p_{mn}^i\phi_{mn} - 2\sum_{m,n} p_{mn}^t\phi_{mn}$$

From here, we will consider each mode separately. Without loss of generality:

$$\left(\widetilde{D}\nabla^4 - m_p\omega^2\right)w_{mn}\phi_{mn} = 2p_{mn}^i\phi_{mn} - 2p_{mn}^t\phi_{mn}$$

First, let us use the biharmonic operator on the mode shape:

$$\nabla^4\phi_{mn} = \frac{d^4}{dx^4}\phi_{mn} + 2\frac{d^2}{dx^2}\frac{d^2}{dy^2}\ \phi_{mn} + \frac{d^4}{dy^4}\ \phi_{mn}$$
$$= \left(\frac{m\pi}{L_x}\right)^4 \phi_{mn} + 2\left(\frac{m\pi}{L_x}\right)^2\left(\frac{n\pi}{L_y}\right)^2 \phi_{mn} + \left(\frac{n\pi}{L_y}\right)^4 \phi_{mn}$$

Plugging this back in we get:

$$\left(\widetilde{D}\left[\left(\frac{m\pi}{L_x}\right)^4 + 2\left(\frac{m\pi}{L_x}\right)^2\left(\frac{n\pi}{L_y}\right)^2 + \left(\frac{n\pi}{L_y}\right)^4\right]\phi_{mn} - m_p\omega^2\phi_{mn}\right)w_{mn} = 2p_{mn}^i\phi_{mn} - 2p_{mn}^t\phi_{mn}$$

We can simplify out the common $\phi_{mn}$ term and expand $\widetilde{D}$ to yield:

$$\left(D(1+j\eta)\left[\left(\frac{m\pi}{L_x}\right)^4 + 2\left(\frac{m\pi}{L_x}\right)^2\left(\frac{n\pi}{L_y}\right)^2 + \left(\frac{n\pi}{L_y}\right)^4\right] - m_p\omega^2\right)w_{mn} = 2p_{mn}^i - 2p_{mn}^t$$

Now, let us make some definitions. First, let us define the modal wavenumber as:

$$k_{mn} = \sqrt{k_m^2 + k_n^2} = \sqrt{\left(\frac{m\pi}{L_x}\right)^2 + \left(\frac{n\pi}{L_y}\right)^2}$$

Next, let us define the modal angular frequency as:

$$\omega_{mn} = \sqrt{\frac{D}{m_p}k_{mn}^2}$$

Then, if we pull out the $m_p$ term and substitute the modal angular frequency we can write:

$$m_p(\omega_{mn}^2(1+j\eta) - \omega^2)w_{mn} = 2p_{mn}^i - 2p_{mn}^t$$

With some rearrangement we get:

$$\frac{m_p}{2}(\omega_{mn}^2 - \omega^2 + j\eta\omega_{mn}^2)w_{mn} = p_{mn}^i - p_{mn}^t$$

We can determine the pressure transmitted from the plate also by considering the normal velocity of the plate and multiplying it by the acoustic impedance of the plate:

$$p_{mn}^t = \sum_{pq} Z_{mn,pq}V_{mn}$$

Note that here, our acoustic modal impedance is defined by two pairs of modes; for completeness, we are describing the effects of intermodal coupling on the pressure. Previous work on the radiation efficiency of submerged plates has shown that the effect of mutual coupling (i.e. $mn \neq pq$) is negligible [6], so we will ignore mutual-impedance and only focus on self-impedance:

$$p_{mn}^t = Z_{mn}V_{mn}$$

Now, note that $V_{mn} = j\omega w_{mn}$ so we can rewrite the modal displacement as:

$$w_{mn} = \frac{V_{mn}}{j\omega}$$

Plugging in our expression for the modal displacement along with our expression for the modal transmitted pressure into the modal equation of motion:

$$\frac{m_p}{2}(\omega_{mn}^2 - \omega^2 + j\eta\omega_{mn}^2)\frac{V_{mn}}{j\omega} = p_{mn}^i - Z_{mn}V_{mn}$$

Now rearranging for $V_{mn}$

$$V_{mn} = \left[\frac{m_p}{2j\omega}(\omega_{mn}^2 - \omega^2 + j\eta\omega_{mn}^2) + Z_{mn}\right]^{-1} p_{mn}^i$$

The modal impedance is a complex term:

$$Z_{mn} = \Theta_{\mathrm{mn}} + j\chi_{mn}$$

Where the real part, $\Theta$, is often referred to as the modal radiation efficiency and the imaginary part, $\chi$, represents the radiation reactance, which is the result of virtual mass loading due to the surrounding fluid. There is not yet a complete solution for both of these terms, but various approximations have been given over the years. The modal radiation efficiency has been solved for all frequencies by Wallace [8] and can be expressed as:

$$\theta_{mn} = \rho_0 c_0 \sigma_{mn}\,,\ \sigma_{mn} = \frac{64k_0^2 L_x L_y}{\pi^6 m^2 n^2}\int_0^{\pi/2}\int_0^{\pi/2}\left\{\frac{\begin{matrix}\cos\\ \sin\end{matrix}\left(\frac{\alpha}{2}\right)\begin{matrix}\cos\\ \sin\end{matrix}\left(\frac{\beta}{2}\right)}{\left[\left(\frac{\alpha}{m\pi}\right)^2-1\right]\left[\left(\frac{\beta}{n\pi}\right)^2-1\right]}\right\}^2 sin\theta d\theta d\psi$$

Where $\alpha = k_0 L_x \sin(\theta)\cos(\psi)$, $\beta = k_0 L_y \sin(\theta)\sin(\psi)$, and the use of $cos$ or $sin$ in the integrand depends on whether $m$ is odd ($\cos(\alpha/2)$) or even ($\sin(\alpha/2)$) and whether $n$ is odd ($\cos(\beta/2)$) or even ($\sin(\beta/2)$).

The radiation reactance is a strong function of frequency. A completely analytical expression over all frequencies has not been determined, but an approximation used by Lomas and Hayek (1977) can be used to good accuracy:

$$\chi_{mn} = \rho_0 c_0 \begin{Bmatrix} k_0/k_{mn} & \frac{k_0}{k_{mn}} \le 1 \\ 0 & \frac{k_0}{k_{mn}} > 1 \end{Bmatrix}$$

This expression essentially takes into account that at high frequencies, the effect of water loading is negligible. To simplify the expressions, we will use the symbolic radiation impedance for the rest of the analysis.

Substituting the components of the modal impedance:

$$V_{mn} = \left[\frac{m_p}{2j\omega}(\omega_{mn}^2 - \omega^2) - \frac{\chi}{j} + \frac{m_p\eta\omega_{mn}^2}{2\omega} + \Theta\right]^{-1} p_{mn}^i$$

$$V_{mn} = \left[\frac{m_p}{2j\omega}\left(\omega_{mn}^2 - \frac{2\omega\chi}{m_p} - \omega^2\right) + \frac{m_p\eta\omega_{mn}^2 + 2\omega\Theta}{2\omega}\right]^{-1} p_{mn}^i$$

$$V_{mn} = \frac{2j\omega}{m_p}\left[\left(\left(\omega_{mn}^2 - \frac{2\omega\chi}{m_p}\right) - \omega^2\right) + j\omega_{mn}^2\left(\eta + \frac{2\omega\Theta}{\mathrm{m_p}\omega_{mn}^2}\right)\right]^{-1} p_{mn}^i$$

Now let us define:

$$(\omega_{mn}^e)^2 = \omega_{mn}^2 - \frac{2\omega\chi}{m_p} \qquad \eta^e = \eta + \frac{2\omega\Theta}{m_p\omega_{mn}^2}$$

We can then write:

$$V_{mn} = \frac{2j\omega}{m_p}[((\omega_{mn}^e)^2 - \omega^2) + j\omega_{mn}^2\eta^e]^{-1} p_{mn}^i$$

Frequently the modal amplitude of the plate velocity is expressed in terms of an admittance:

$$V_{mn} = Y_{mn} p_{mn}^i$$

Where the modal admittance is:

$$Y_{mn} = \frac{2j\omega}{m_p}[((\omega_{mn}^e)^2 - \omega^2) + j\omega_{mn}^2\eta^e]^{-1}$$

Recall the formula for acoustic power is given by:

$$\Pi = \frac{1}{2} Re\{\int_S Pv^* ds\}$$

Where $S$ is the area being integrated over. The incident power can be given by :

$$\Pi_i = \frac{L_x L_y |p_i|^2}{2\rho_0 c_0}$$

To solve for the transmitted power, we can rewrite this expression in terms of modes:

$$\Pi_{\mathrm{t}} = \frac{1}{2} Re\{\sum_{m,n} Z_{mn} V_{mn} V_{mn}^*\} = \frac{1}{2} Re\{\sum_{m,n} Z_{mn} |V_{mn}|^2\} = \frac{1}{2} Re\{\sum_{m,n} Z_{mn} \, |Y_{mn}|^2 \left|p_{mn}^i\right|^2\}$$

The power transmission coefficient can then be given by:

$$|T|^2 = \frac{\Pi_t}{\Pi_i}$$
